# Intuitive sensorimotor decisions under risk take Newtonian physics into account

**DOI:** 10.1101/2025.01.07.631692

**Authors:** Fabian Tatai, Dominik Straub, Constantin A. Rothkopf

**Affiliations:** Centre for Cognitive Science, Technical University Darmstadt, 64283 Darmstadt, Germany; Institute of Psychology, Technical University Darmstadt, 64283 Darmstadt, Germany; Hessian Center for Artificial Intelligence, Darmstadt, Germany

**Keywords:** decision-making, intuitive physics, sensorimotor control, utility theory, object interaction, rational analysis, mixed reality

## Abstract

The success of our interactions with objects in natural tasks depends on generating motor actions, which are subject to sensorimotor variability, the objects’ physical properties, which are governed by the laws of kinematics, and the costs and benefits associated with the actions’ outcomes. Therefore, such interactions jointly involve sensorimotor control, intuitive physics, and decision-making under risk. Here, we devised a mixed reality experiment, which allows investigating human behavior involving the interaction of all three of these cognitive faculties. Participants slid pucks to target areas associated with monetary rewards and losses within an immersive, naturalistic virtual environment. In this task, signal-dependent variability inherent in motor control interacts non-linearly with the physical relationships governing objects’ motion under friction. By systematically testing generative models of behavior incorporating different assumptions about how participants may generate the slides, we find decisive evidence that participants’ decision under risk readily took their individual motor variability and Newtonian physics into account to gain monetary rewards without relying on trial by trial learning.

**Author summary:** Humans are exceptionally proficient at manipulating objects in natural tasks. This skill requires integrating three cognitive faculties: 1. Sensorimotor-control, subject to inherent uncertainty and variability; 2. Intuitive physical reasoning as object manipulations are subject to physical dynamics; 3. Economic decision-making when actions have outcomes with monetary consequences. Here, we show that in a naturalistic, immersive object manipulation task, human participants plan their movements by taking into account how their sensorimotor variability interacts with the physical dynamics of object manipulation to optimize economic outcomes. Therefore, our findings suggest that in everyday naturalistic interactions with the world, humans can integrate these three cognitive faculties.

## Introduction

Imagine a bartender skillfully sliding a drink across a bar counter into the hands of a customer. The success of the sliding action depends on three factors: the generated sensorimotor action, which is subject to motor variability, the sliding movement of the glass, which is governed by the kinematic laws of physics, and the costs and benefits associated with the potential outcomes. Accordingly, from the perspective of the bartender, selecting a puck’s release velocity or an aim point on the counter involves three cognitive faculties: sensorimotor control [1–4], intuitive physics [5–8], and value-based decision-making under risk [9–12]. Therefore, such a sliding task allows investigating the interaction of these three cognitive faculties in a setting that is much more naturalistic [13–15] compared to many laboratory tasks that are commonly used to study the three involved cognitive faculties in isolation.

While classically decision-making under risk has been studied in economic domains using repeated choices between lotteries [10], more recently formally equivalent sensorimotor decisions [16, 17] have been investigated [18–23], in which the probabilistic outcome arises due to the stochasticity in the sensorimotor system [24]. Statistical decision theory (SDT) prescribes that the bartender should choose to aim at a location on the counter, which maximizes expected utility 𝔼 [*u*(*x*)] [25, 26] across all possible final locations *x*:

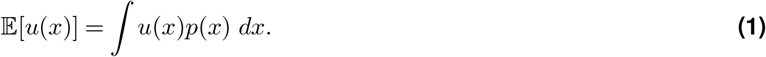

Here, the term *p*(*x*) is the probability of the glass coming to a rest at location *x* determined by the variability in the bartender’s sliding. In perceptual and motor domains, many studies have shown that humans are well calibrated to their perceptual and motor variability and act according to SDT [27–29]. The utility *u*(*x*) in the above equation captures the costs and benefits of each possible final location *x* of the drink. The bartender would like to avoid the drink from spilling over the customer, but still end up reasonably close. If the utility of reaching location *x* is identical to the monetary payoffs *v*(*x*) obtained from reaching *x*, then *u*(*x*) = *v*(*x*). However, human weighting of payoffs is often better described with subjective utility functions *u*(*x*) [25, 26]. Specifically, in economic domains, humans have long been known not to maximize expected utility [11, 12], requiring further assumptions including loss-aversion to describe human decisions better [10]. By contrast, in sensorimotor tasks that are equivalent to such economic decisions, many studies have shown that human movement choices are well described by SDT given participants’ individual motor variability [16, 21, 23]. Still, some studies have pointed out that accuracy depends on factors such as training [21], target symmetry [30], and outcome ratio [31], and some studies found that including a non-linear utility weighting *u*(*x*) can better describe human sensorimotor choices under risk [19].

What is missing in these studies, is that sensorimotor interactions in everyday tasks are governed by the laws of physics, which adds a layer of complexity compared to commonly studied sensorimotor actions, e.g. very short pointing or reaching movements, which are often additionally constrained to lie in a single movement dimension by a manipulandum. In the bartender example, the final location of the glass is determined by the initial velocity due to the force produced by the bartender and additionally by the friction of the counter. This situation can be described using the fundamental kinematic equations [32] corresponding to Coulomb friction [33], where an object’s final velocity *v* and displacement *x* depend on its initial velocity *v*_0_ and friction’s deceleration *a*,

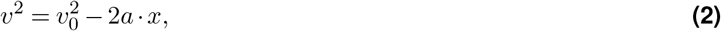

which results in the final position of the puck being a non-linear function of the velocity

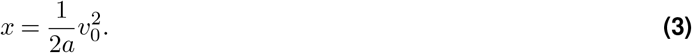

This non-linear function transforms the distribution of intial velocities *v*_0_ caused by sensorimotor variability in the bartender’s sliding movements to the distribution of the final positions *x* of the glass on the counter. Therefore, the question arises whether humans’ sensorimotor decisions under risk take the way that sensorimotor variability interacts with physical kinematics into account. Early research on intuitive physics used forced choice, explicit reasoning tasks, showing pervasive failures of people’s predictions of object’s kinematic trajectories [5, 34], which lead to the hypothesis that people employ simple heuristics [35]. However, the accuracy of predictions depend on familiarity [36], context [37], or the specific task [38]. More recently, people’s intuitive physical reasoning has been found to be consistent with Newton’s laws when accounting for perceptual and internal model uncertainty [6], leading to the hypothesis that humans posses an internal noisy physics engine [7, 39, 40]. However, only recently intuitive physical understanding has been investigated with sensorimotor actions instead of forced choice judgements, showing that people use inferred physical properties of objects in subsequent sensorimotor interactions [41] and that they can use correct kinematic priors [42].

Here, we examine whether human sensorimotor decisions under risk when interacting with a real physical object reflect the impact of kinematics on the variability of movement outcomes. In an immersive virtual reality (VR) setup, participants slid real hockey pucks at targets, yielding monetary gains and losses. Because of movement variability, specifically signal-dependent variability by which the standard deviation of muscle force grows linearly with its mean [43–45], the outcome of the motor action is stochastic. However, this task does not only require naturalistic sensorimotor control, but includes the additional complexity of manipulating an external object. The variability of subjects’ motor actions interacts with the kinematic laws of physics [32], which scale non-linearly by additionally increasing variability for longer slides. Finally, taking the economic consequences of the actions into account, SDT predicts that subjects should undershoot the target more, the higher overshooting is penalized. But not only that, a rational actor should also take the physics of object sliding into account, and therefore should not only undershoot more with higher prospective penalty but even more for more distant targets, because of the increased signal-dependent variability. Finally, we allow for a subjective utility scaling of outcomes [46], as this has been shown better to describe sensorimotor decisions in some previous studies [19].

We carefully constructed a series of Bayesian generative models of sliding actions to examine the contribution of the three cognitive faculties important in our task: sensorimotor control, intuitive physical understanding and weighting of prospective outcomes. Using Bayesian model comparison, we find decisive evidence in favor of all models according to which participants’ sensorimotor decisions under risk readily took the kinematic laws of physics into account without relying on error learning across trials. The single model best accounting for participants’ behavior incorporates an individual’s signal-dependent motor variability, the non-linear scaling prescribed by the Newtonian kinematic laws, and a utility function describing participants’ subjective weighting of penalties.

## Results

Subjects slid real standard hockey pucks at targets to maximize an in-game score, proportionally paid out as a monetary incentive at the end of the experiment. The puck was tracked by a motion capturing system and subjects saw it rendered in a head-mounted VR display (Fig. 1a, SI Apparatus, SI Movie S1). After subjects released the puck, its velocity was taken as the initial velocity *v*_0_ to simulate the puck’s displacement *x* via motion with constant deceleration (see Materials and Methods). In this setup, subjects can naturalistically interact with the puck on the table’s surface, enabling a haptic experience of the friction between the table and the puck. At the same time, it allows efficiency in the data collection, measurements of motor actions, and control over the physics. In previous work, we have shown that even without receiving visual feedback about puck trajectories, this embodied setup leads to intuitive behavior in line with the physics of friction in subjects [42].

**Figure 1.**
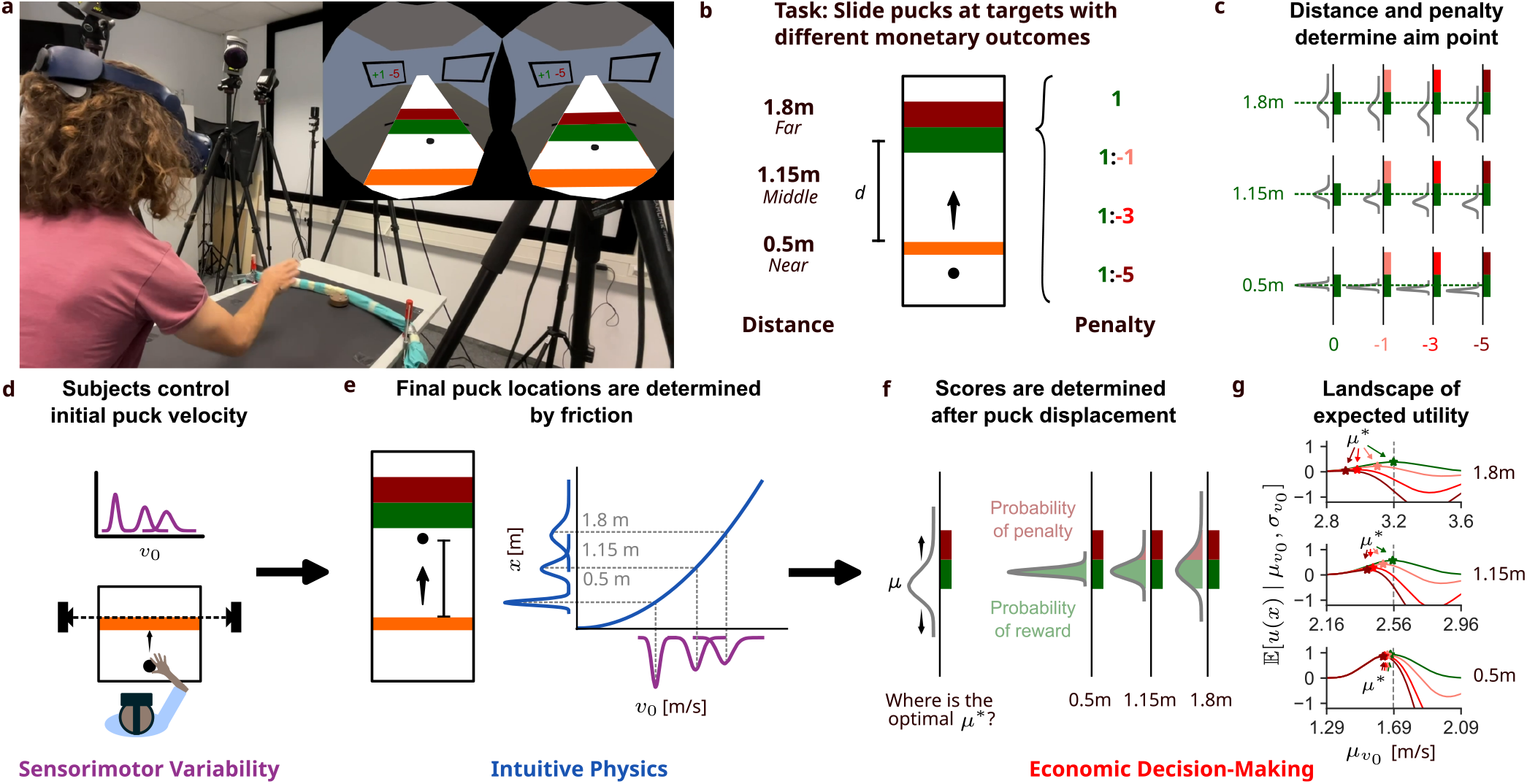
Experimental setup and conceptual overview. **a** The setup: Subjects interacted with a real standard hockey puck and released it into a fabric barrier. The puck’s initial velocity was captured by a motion-capturing system, and its simulated movement in the virtual environment was displayed to the subject via a head-mounted display. (The person shown is the first-author of the manuscript.) **b** The task: Human subjects (*N* = 20) slid pucks at targets, which yielded different scores, forming the basis of a performance-based bonus payment. Each condition was presented once per session on two consecutive days, always in blocks of 30 trials. **c** Statistical decision theory predicts that subjects should undershoot to avoid hitting the penalized area. The higher the prospective penalty, the shorter they should aim. As variability and with that the probability of unwantedly triggering a penalty increases with distance, subjects should undershoot more if the targets are farther away. **d** Generating velocities is subject to signal-dependent variability: Larger distances lead to higher variability. **e** The quadratic mapping from initial puck velocity to final puck location due to kinematics further increases the variability (See SI Kinematics Equations for details). **f** When deciding where to aim (choosing *µ*_*v*_), subjects should consider this increase in variability, as the probability of hitting a target decreases while the probability of being penalized increases with distance. **g** The expected utility of different aim points and median velocities *µ*_*v*_ determine the optimal aim point *µ*^*^ for a target configuration.

In our task, subjects were instructed to hit the green target area (Fig. 1b), which yielded a score of 1 point. In most conditions, the targets were configured such that overshooting the green target and hitting the adjacent red penalty area resulted in a negative score of −1, −3, or −5, depending on the experimental condition. In one target configuration, the red area was not present, and overshooting was not penalized. Ending up anywhere else on the table amounted to a score of 0. Additionally, the center of the target was placed at three different distances: 0.5*m*, 1.15*m*, or 1.8*m*. The total of 12 target conditions (4 penalties, 3 distances) were completed in blocks of 30 trials. Before encountering the penalized conditions for the first time, subjects trained only on the green target to familiarize themselves with the friction of the table’s surface (see SI Task and Procedure for more details).

Essentially, the subjects’ task was to generate an initial puck velocity *v*_0_ such that the puck would hit the target but avoid the penalty area (Fig. 1c). Generating velocities is subject to signal-dependent variability [43–45], as higher velocities evoke higher variability in the subjects’ motor systems (Fig. 1d). This can be well described with a log-normal distribution over velocities parameterized by the median 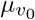 of this distribution. The initial velocity *v*_0_ is transformed by the kinematics of motion with constant deceleration, resulting in a final puck position that is a quadratic function of the initial velocity (Fig. 1e). The consequence is that hitting a target of the same size is much more probable when it is closer rather than farther away (Fig. 1e). However, as some conditions also include a penalty area, maximizing the outcome not only needs to account for motor variability and the kinematics of sliding, but also consider that overshooting the target might result in a penalty (Fig. 1f). Accordingly, there is an optimal initial velocity 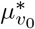 that maximizes expected utility (Fig. 1g).

### Participants immediately undershoot more with higher penalty and larger distance

For an initial statistical analysis of the influence of the target conditions on subject behavior (Fig. 2a), we fit Bayesian linear mixed-effects models (see SI Mixed Effects Models for details). First, we confirmed (see SI Fig. S1) that the pucks’ initial velocities showed signal-dependent variability (see Fig. 2b) and that friction increased the corresponding final location variability across target distances (see Fig. 2c). Accordingly, SDT makes the following two main qualitative predictions how subjects can maximize their performance, (Fig. 1c): 1. Subjects should slide shorter the higher the prospective penalty, because this reduces the probability of overshooting the target, i.e., hitting the penalized area. 2. Subjects should slide shorter, the larger the target distance, because the larger variability for longer slides increases the probability of unwantedly hitting the penalized area.

**Figure 2.**
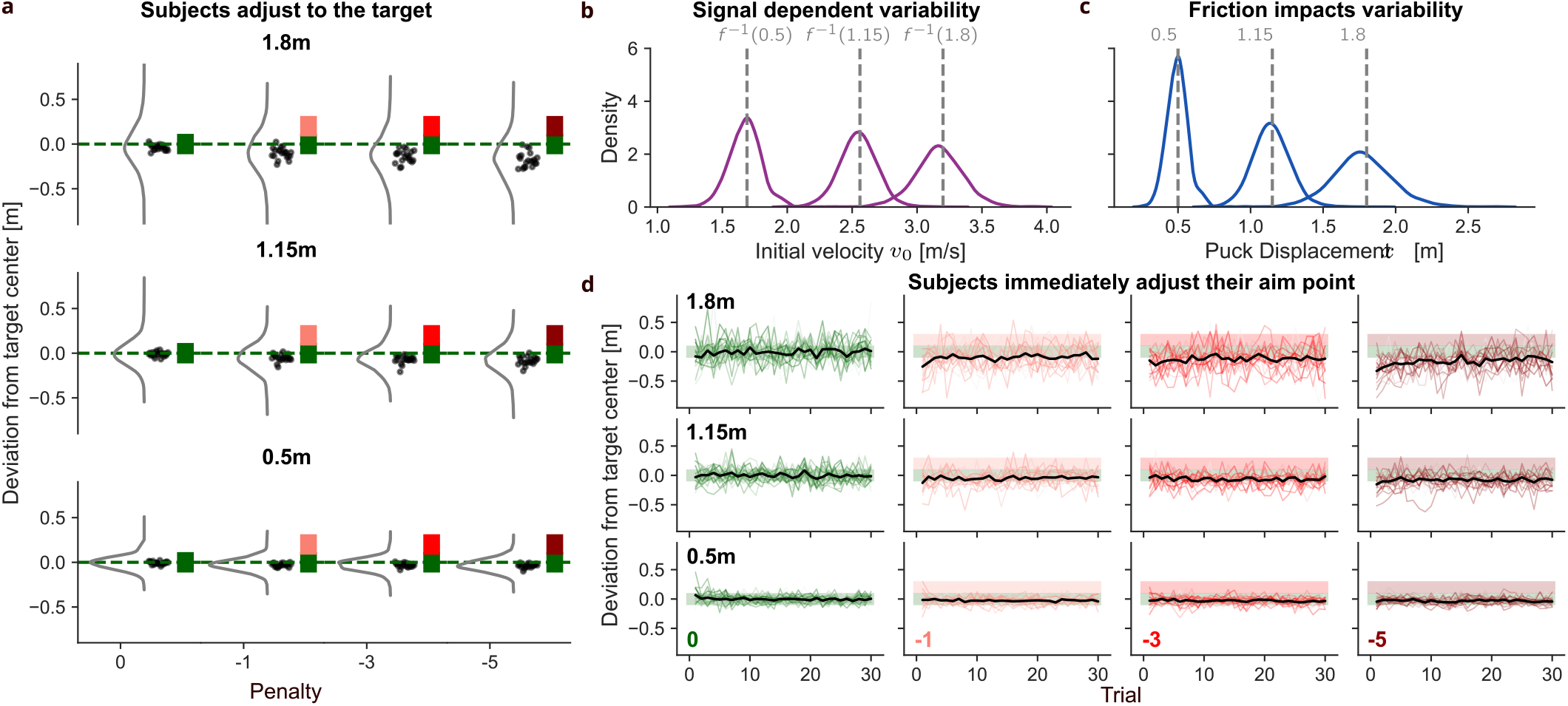
Behavioral data. **a** Final puck location relative to the target center per condition across all subjects (lines) and median per subject (dots). We find that both manipulations (penalty and target distance) have a significant effect on sliding behavior, i.e., subjects undershoot more the higher the distance and the higher prospective penalty. **b** Initial puck velocity in the condition with 0 penalty across the target distances. Variability significantly increases with target distance. **c** The kinematics of friction non-linearly scale and further increase variability. **d** Progression of slides over the course of the first session per condition. Colored lines correspond to one subject, and black lines are the means across subjects. Subjects immediately undershoot the target when penalty is present. This suggests that no learning is necessary as subject behavior transfers to novel target configurations.

To test this, we fit a mixed effects model (see SI Statistical Analysis with Mixed Effects Models), which confirms that subjects undershoot more with higher distance, but also with higher penalty (see SI S2). Importantly, we find that penalty and target distance interact, as at target distance 0.5*m* the effect of penalty −5 is small (*β*_0,0.5*m*_ − *β*_−5,0.5*m*_ = −0.04, 95% CI [0.01, −0.05]) and the largest distance 1.8*m* the effect is much bigger (*β*_0,1.8*m*_ − *β*_−5,1.8*m*_ = −0.18, 95% CI [−0.21, −0.16]). At the highest penalty and distance, participants even aimed considerably outside the target (median: 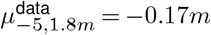, relative lower target area bound *A*_*l*_ − 1.8*m* = −0.1*m*). This interaction is qualitatively in accordance with kinematic’s influence in SDT, as the much lower variability at 0.5*m* should cause much smaller effects of penalty.

To exclude the possibility that subjects adjusted their sliding over trials using an error learning strategy, we hypothesized that subjects could readily avoid overshooting implying an internal representation of the kinematic’s impact on the slide variability. To test this, we modeled the distance to the target with a term for trials in a block and allowed it to differ across conditions. Across the different conditions, we find a significant, systematic effect only at the farthest target distance 1.8*m*, see also Fig. 2d. However, subjects start out with more undershoot and then successively start sliding closer to the target (e.g. penalty −5: *β*_trial,−5,1.8_ = 0.11, 95% CI [0.08, 0.15]). Importantly, this effect is also present in the no penalty condition at 1.8*m* (*β*_trial,0,1.8_ = 0.04, 95% CI [0.00, 0.07]) (see SI Fig. S2 for all conditions). This suggests that the effects of the different target configurations on sliding behavior is stronger at the start of a block at distance 1.8*m*. Another confirmation of this finding comes from re-fitting the mixed effects model on only the first trial that participants completed in each condition (see SI Figure S3). Therefore, a purely error-based learning strategy seems highly unlikely, as subjects immediately choose an appropriate strategy.

### Participants take into account how their individual variability interacts with the kinematics

To describe quantitatively how closely subjects’ behavior agrees with SDT, we devised a series of Bayesian generative models of sliding behavior incorporating different assumptions (Fig. 3a). The *Physics* model is an instantiation of a rational actor based on SDT which plans the sliding movement using the correct sensorimotor variability, the correct kinematics of the task, and uses the outcomes as utilities. We assume that each participant *p* generates initial puck velocities with their own personal variability *σ*_*p*_ according to Equation 7. This in turn determines where a participant should aim 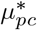 in condition *c* under the assumption that expected utility is maximized with respect to individuals’ variability of final puck location determined by *σ*_*p*_ and the kinematics of the task (Equation 6; see SI Models for details). Fig. 3b shows that subjects are mostly in line with predictions of our task analysis, as empirical aim points 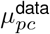 (medians *µ* for participant *p* in condition *c*) fall shorter the higher the distance and the higher the prospective penalty.

**Figure 3.**
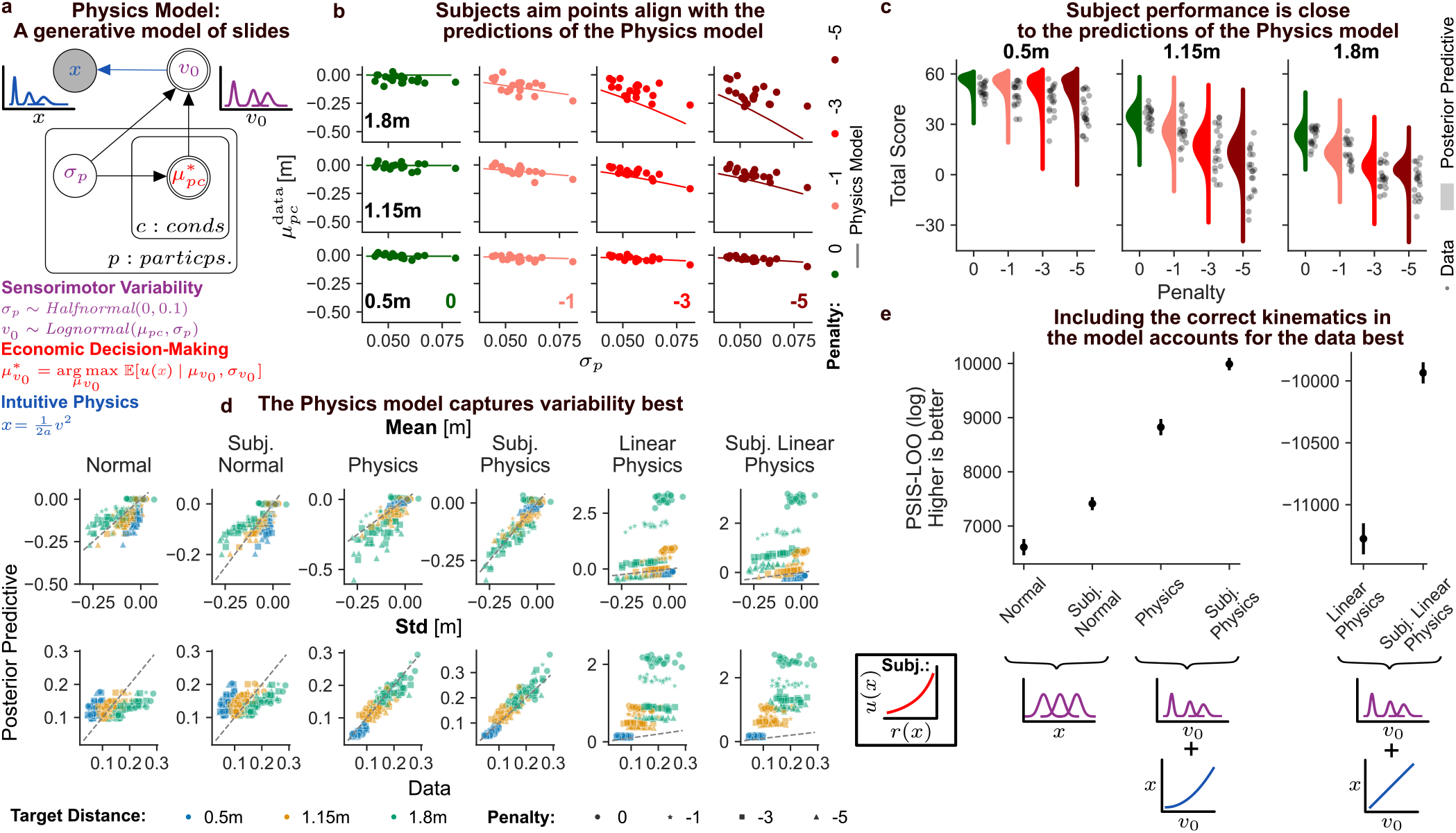
**a** The *Physics* model is based on statistical decision theory, includes signal dependent variability of initial puck velocity and accounts for sliding kinematics. **b** The participants’ variability *σ*_*p*_ as estimated by the *Physics* model compared to the participants’ median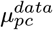. Participants’ behavior is close to the predictions, as undershoot increases with variability, target distance and prospective penalty. **c** Subject score per condition (60 trials) compared to the score of the posterior predictive of the *Physics* model corresponding to playing 10000 times. Even in the high penalty condition at a larger distance, where subjects’ aim is less careful than predicted by the model, performance is still quite close. Using posterior predicitives **d** and approximated leave-one-out cross validation scores (*PSIS-LOO*) **e** we compare different models’ predictions of our data. Assuming no signal dependent variability (*Normal* model), or internal linear approximation of the kinematics (*Linear Physics*) misrepresents the data. Therefore, including the correct kinematics (*Physics*) seems to explain the data best. All models, benefit when a subjective utility function is added.

As we infer subjects’ variability individually, we can test another central prediction of SDT. Not all subjects are equally accurate at sliding pucks, i.e., some have higher variability than others. Similar to the increase of variability with distance, this predicts that rational actors whose slides are more variable should undershoot more than those with more precise sliding. To test this, we add the individual variability *σ*_*p*_ inferred with the *Physics* model as an additional predictor to the mixed-effects model predicting slide deviation from the target center. We find that subjects with higher variability undershoot the target more 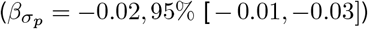 (see SI Fig. S2 for effects per condition). This provides further evidence that subjects used their individual motor variability in generating the sliding action.

While participants’ sliding is well described in the −1 penalty condition, at larger distances and higher penalties (−3 and −5), participants did not undershoot as much as predicted by the *Physics* model. However, a comparison between subjects’ scores in the task with the expected outcome as predicted by the model (Fig. 3c) gives an indication why this might be the case. Generally, subjects did not perform much worse, and some were even on par with the rational actor, which has a low expected utility for larger distances and a high prospective penalty. For example, at 1.8*m* and penalty −5 the total penalty on average was −8 cents compared to the optimal strategy of 2 cents at the final pay out. Considering that subjects earned a bonus of 5.77 € on average, this difference only amounts to about 2%. This can be explained with the relatively flat surface of the expected utility, see Fig. 1f, implying that aim points slightly closer to the target than the optimum yield only slightly worse expected outcomes, which are almost indistinguishable from the optimal strategy.

### Participants’ behavior is best described by including Newtonian kinematics

To investigate whether participants’ sliding behavior reflected the correct physical kinematics, we implemented a series of models incorporating different assumptions about sensorimotor variability, the kinematics of sliding, and the subjective utility of outcomes. The reasoning for the choice of models was to systematically compare models incorporating different assumptions from previous studies on sensorimotor decisions under risk with either the correct Newtonian physics or, instead, a linear heuristic in the planning of sliding actions. This linear heuristics was implemented as a possible arbitrary linear scaling of the final distance relative to participants’ sliding action. In all models, the aim points within conditions are chosen to maximize utility, but the assumptions about the underlying variability and physics change. Additionally, we compared these models to a baseline model proposed by Trommershäuser, Maloney, and Landy [18] in a sensorimotor pointing task under risk. Table 1 lists all considered models together with their respective assumptions, and further details are provided in the Methods section. To evaluate the different models, we observe how well posterior predictive distributions of different models account for the empirical data (Fig. 3d) and carried out model comparison by using Pareto-smoothed importance sampling (PSIS-LOO) as an approximation to leave-one-out cross-validation [47]. Importantly, PSIS-LOO is an estimate of the out-of-sample prediction accuracy and, therefore, takes a model’s complexity into account.

**Table 1.**
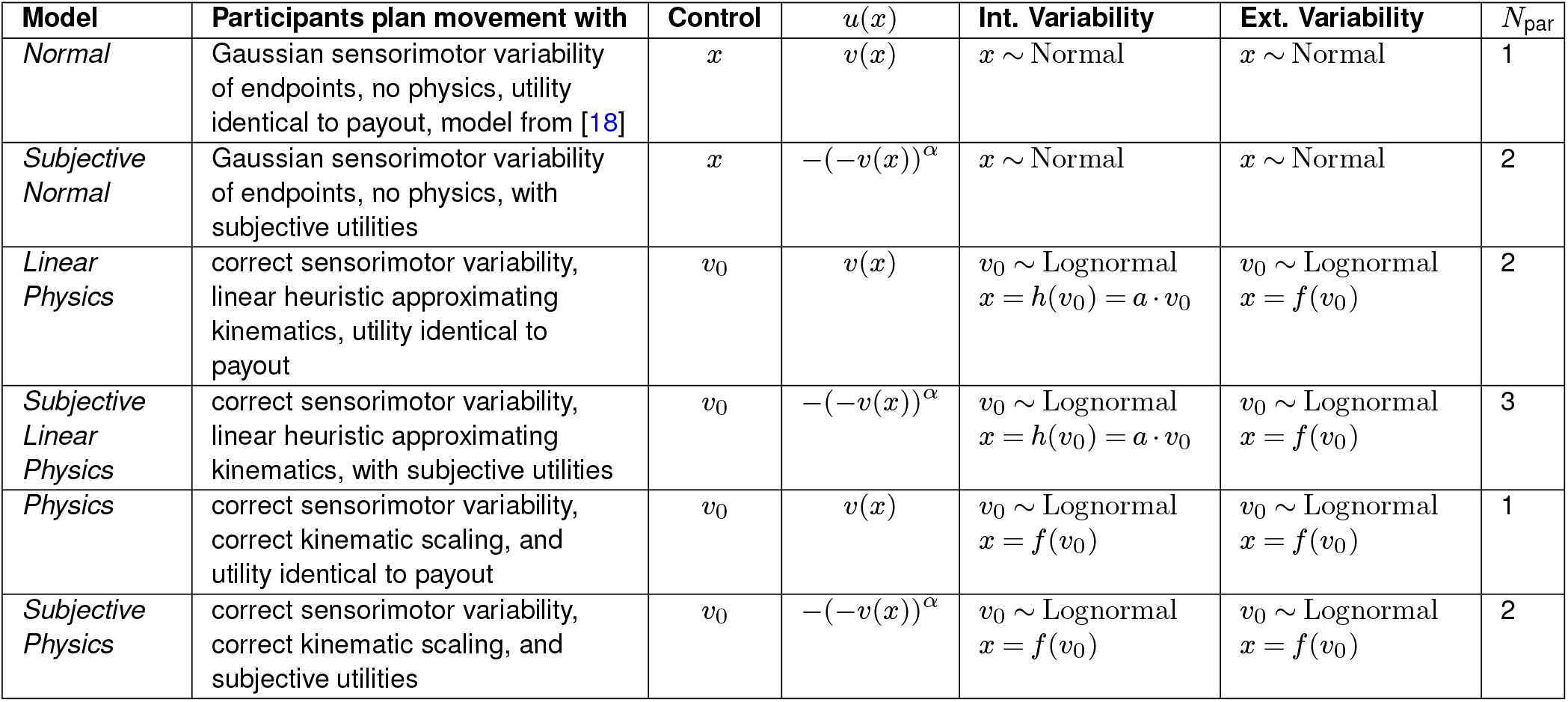
Overview over models and their assumptions. **Model**: model name, **Control**: the variable that the subject is assumed to control, *u*(*x*) the subject’s utility function (veridical: *u*(*x*) = *v*(*x*) or subjective see Equation 12). **Internal Variability**: the internal variability that the subject assumes to determine optimal aim points, **Ext. Variability**: the external variability underlying the generative model of slides, *N*_par_: the number of free parameters per subject.

| Model | Participants plan movement with | Control | $u(x)$ | Int. Variability | Ext. Variability | $N_{\text{par}}$ |
| --- | --- | --- | --- | --- | --- | --- |
| *Normal* | Gaussian sensorimotor variability of endpoints, no physics, utility identical to payout, model from [18] | $x$ | $v(x)$ | $x \sim \text{Normal}$ | $x \sim \text{Normal}$ | 1 |
| *Subjective Normal* | Gaussian sensorimotor variability of endpoints, no physics, with subjective utilities | $x$ | $-(-v(x))^\alpha$ | $x \sim \text{Normal}$ | $x \sim \text{Normal}$ | 2 |
| *Linear Physics* | correct sensorimotor variability, linear heuristic approximating kinematics, utility identical to payout | $v_0$ | $v(x)$ | $v_0 \sim \text{Lognormal}$<br>$x = h(v_0) = a \cdot v_0$ | $v_0 \sim \text{Lognormal}$<br>$x = f(v_0)$ | 2 |
| *Subjective Linear Physics* | correct sensorimotor variability, linear heuristic approximating kinematics, with subjective utilities | $v_0$ | $-(-v(x))^\alpha$ | $v_0 \sim \text{Lognormal}$<br>$x = h(v_0) = a \cdot v_0$ | $v_0 \sim \text{Lognormal}$<br>$x = f(v_0)$ | 3 |
| *Physics* | correct sensorimotor variability, correct kinematic scaling, and utility identical to payout | $v_0$ | $v(x)$ | $v_0 \sim \text{Lognormal}$<br>$x = f(v_0)$ | $v_0 \sim \text{Lognormal}$<br>$x = f(v_0)$ | 1 |
| *Subjective Physics* | correct sensorimotor variability, correct kinematic scaling, and subjective utilities | $v_0$ | $-(-v(x))^\alpha$ | $v_0 \sim \text{Lognormal}$<br>$x = f(v_0)$ | $v_0 \sim \text{Lognormal}$<br>$x = f(v_0)$ | 2 |

We start with models that work directly in position space, i.e. do not consider the physical relationship between the generated velocities and the final puck positions. Therefore, decisions in this model are in the space of final puck positions instead of velocities. These models are equivalent to the models by Trommershäuser, Maloney, and Landy [18], as they do not account for signal-dependent motor variability. As such, these models serve as a baseline to check whether assumptions of the model by Trommershäuser et al. [18] are sufficient to capture the sliding behavior under the influence of Newtonian kinematics. This model assumes a normal distribution of final puck position *x* with the same *σ* for each distance over the slide outcomes (*Normal* model). The posterior predictive indicates that the model misrepresents the data, as it does not correctly describe the mean and standard deviation across conditions. We find that this model performs far worse than the *Physics* model (*d*_LOO_ = 2214.89, *se*_*d*_ = 123.54). This does not come as a surprise, as the *Normal* model can not account for the known phenomenon of signal-dependent variability. Therefore, it seems strictly necessary to include this aspect in a model. In accordance with this, the *Physics* models’ posterior predictive (see Fig. S4) shows that it characterizes the data much better. This is evidence that signal-dependent variability is needed to account for subjects’ behavior.

However, variability in the *Physics* model increases because of the combination of signal-dependent variability in generating velocities and the physical relationship connecting velocities to slide outcomes. While it seems reasonable that subjects are adapted to the increase in variability of initial puck velocities, as they generate those with their own actions [1], it is unclear whether subjects can account for the effects of the table’s friction. Therefore, we have constructed a model *Linear Physics*, assuming that subjects employ a linear heuristic *h* instead of the actual physical relationship predicting the relationship between initial velocity *v*_0_ and the final puck position *x*:

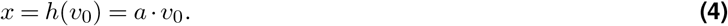

For each participant *p* we treat the slope *a*_*p*_ as a free parameter of the model. With this model, we test the hypothesis whether it would suffice for the subjects to simply approximate the physics of the task, rather than veridically representing it. By treating the slope as a free parameter, we infer the subject’s internal approximation as the one that most likely produced our data. Note that while the *Linear Physics* model assumes that the subjects have a subjective internal linear approximation of the physics, we model the actual generative process of the slides according to the correct physics. Thus, this model explicitly assumes that human participants have a false belief about the motion of an object they are interacting with, a phenomenon for which previous research has found quantitative evidence [48]. The posterior predictive (see Fig. 3d) of the *Linear Physics* model shows that this assumption can not at all account for the subjects’ data. Especially for far distances, the linear approximation leads to a big mismatch of the real friction. The subjects would overestimate the velocity needed to hit the target, which, because of the quadratic relationship governing the real friction, will lead to large overshoots of the target (see SI Fig. S4). It even performs substantially worse than the *Normal* model (*d*_LOO_ = 17885.68, *se*_*d*_ = 161.95) giving a further indication that its core assumptions are wrong. Therefore, signal-dependent variability in the initial velocities can not solely explain subject behavior, and subjects’ planning involves the correct physical relationship.

The *Physics* model predicts larger undershoots than those observed in the empirical data in the −5 condition (see Fig 3 d). For the farthest distance, this difference amounts to 0.14*m* on average when comparing the posterior predictive mean with the data mean. As previous research has shown that individual subjects may scale outcomes to monetary payouts differently in economic and sensorimotor decisions under risk [19], we extended the previous models by including a subjective utility function for prospective penalty *u*(*x*) = −(−*r*(*x*))*α* for *r*(*x*) *<* 0 [46] for each participant *α*_*p*_, instead of assuming *u*(*x*) = *v*(*x*) like in the plain version of the models. Out of the subjective models, we find that the *Subj. Physics* model performs best and fits the data very closely. This leads to a significant improvement over the plain *Physics* model (*d*_LOO_ = 1164.73, *se*_*d*_ = 51.75) It is noteworthy that even the plain *Physics* model explains the data better than the *Subj. Normal* model (*d*_LOO_ = 1411.97, *se*_*d*_ = 120.05) suggesting once more that accounting for the increase in variability is crucial for modeling the subject data. Furthermore, one could assume that a combination of a sensible subjective utility function combined with an approximation of physics could explain the data, as the two functions offer considerable freedom in expressing how actions and actions outcomes relate (3 free parameters per subject vs. 1 in the plain *Physics* model). This would suggest that subjects would not be calibrated to the physics. However, even though the *Subj. Linear* model is an improvement over its plain version (*d*_LOO_ = 1341.09, *se*_*d*_ = 45.99), it still performs very badly at describing the data, clearly ruling out this hypothesis. The posterior estimates of *α*_*p*_ for the *Physics* model and the corresponding subjective penalty functions are shown in SI Fig. S6, showing a general lowered sensitivity toward penalty (*α*_*p*_ mean: 0.28) with considerable inter-subject variability (*α*_*p*_ min: 0.02, max: 1.02, std: 0.25).

## Discussion

To be successful in naturalistic everyday interactions with objects such as in tool use, food preparation, or sports, people need to integrate motor control, intuitive physical understanding, and weighting prospective outcomes. Previous studies have separately investigated intuitive physical reasoning [5–8], economic decision-making [9–12] or sensorimotor control [1–4]. The work in [16, 19, 49] compellingly integrates sensorimotor control with economic decision-making, and intuitive physics in combination with sensorimotor-control has also been studied [41, 50, 51]. Here, we devised a mixed reality experiment in which participants slid real pucks to targets for monetary outcomes, which allows studying the interaction of all three cognitive faculties in a naturalistic setting. Thus, this task allows investigating the much richer interaction of the three involved cognitive faculties compared to many laboratory tasks used to study these cognitive faculties separately. To understand participants’ behavior, we implemented a series of probabilistic generative models, which allowed us to dissect the contribution of the three cognitive faculties. The models explain sliding behavior by incorporating different assumptions that have been suggested in previous literature. These models include normally distributed versus signal-dependent log-normal motor variability, models incorporating the correct physical kinematics versus models without a notion of kinematics, and a neutral versus subjective scaling of monetary outcomes. Bayesian model comparison showed that the empirical data was explained best by participants taking their individual signal-dependent motor variability [43–45] and the kinematic laws of physics [32] into account and integrating the stochastic movement outcomes with an individual utility function [19, 46]. Importantly, all pairwise comparisons between the different models, including or excluding the physical nonlinear scaling of initial velocities to final puck distances, decisively favored the conclusion that participants’ sensorimotor decisions under risk incorporated Newtonian physics.

Our results also point towards the computations integrating these three cognitive faculties in the brain. Previous studies have identified brain networks that are preferentially engaged in physical reasoning, leading to the notion of a “physics engine” in the brain [40, 52], in accordance with computational models [7, 8]. Other work has focused on regions preferentially involved in economic decision-making [12, 53] or with integrating sensorimotor control [23]. The regions associated with the “physics engine” overlap with those involved in action planning [54], suggesting that representations involved in intuitive physical reasoning may not be fully abstract and disembodied but instead, at least in part, closely integrated with the afforded actions and potentially movement planning. Our behavioral and modeling results support the idea that intuitive physics can not only be integrated with judgments and predictions [52, 54, 55], but also with action planning and sensorimotor control [40, 54, 56]. This is also in line with evidence that physical properties such as mass are represented along the ventral visual pathway [57] and primary motor cortex [58]. However, most of these results are based on passive observations of physical stimuli or mental simulation of physical interactions, whereas the present study investigates intuitive sensorimotor decisions under risk in a naturalistic task involving physical kinematics. The present results of participants’ ability to additionally take the monetary outcomes of their actions into account imply that action planning in physical scenarios can include computations of valuation.

An important additional implication of the present work is the evidence that the involved cognitive systems are sensitive to individual motor uncertainties in interaction with the kinematics and monetary outcomes. Specifically, in our experiment, subjects’ behavior was sensitive to the uncertainty in prospective outcomes, suggesting that intuitive physics can interact with economic decision-making in sensorimotor tasks. Importantly, this integration of uncertainty with physics and outcomes suggests that the outcomes of sensorimotor actions might be represented in a probabilistic fashion, the implementation of which is a topic of ongoing debate [59]. Future studies should investigate how these three cognitive faculties interact and are integrated in the brain when guiding sensorimotor behavior to shed light on humans’ formidable capabilities in manipulating objects in naturalistic everyday tasks [60]. To this end, mixed reality setups, as in the present study, allow studying naturalistic sensorimotor tasks in physical scenarios with the benefit of additional experimental control, which makes it possible to quantitatively compare behavior against theory-driven generative models of behavior.

## Methods

### Participants

20 undergraduate students at the Technical University of Darmstadt, who were naive with regard to the goals of the study, participated either for course credit or an hourly compensation of 10€. Additionally, all participants were awarded a bonus payment of 3 − 8€ proportional to their cumulative score earned during the experiment. The experiment was approved by the local Ethics Committee and all participants gave informed consent.

### Procedure

Subjects participated in the task on two consecutive days. On day one, subjects began with non-scored training with only the target area: 100 trials at target distances between 0.2 − 2m in order to learn the simulated friction and two blocks of 30 trials of each distance used in our experiment. Trials were blocked to avoid sequential effects such as sensorimotor regression to the mean [61] and to ensure enough familiarization [62]. Then, the scored phase followed and subjects encountered the penalty area for the first time. This phase consisted of 12 conditions (3 distances, 4 penalty conditions), each completed in blocks of 30 consecutive trials. The order of the blocks was randomized for each subject. This procedure allowed us to investigate how well subjects understand the kinematics of sliding pucks, because they had to transfer from the training to novel cost functions in the scored phase [63]. On the second day, subjects were only given 10 training trials per distance, and then the scored phase was repeated.

### Apparatus

In our experimental setup (SI Movie S1), participants were placed in a virtual representation of the room they were standing in, rendered through the *HTC Vive Eye Pro* and the *Unity* game engine. They were tasked to slide a real standard hockey puck, which was tracked by a passive motion capture system (6 Cameras operating with 150*Hz, Qualysis Oqus 500+ and 510+*), across a table. The puck was modified with weights such that its mass amounted to 250*g*, and equipped with felt at the bottom to enhance its sliding properties. To track the puck, four markers were placed on top of it. Thus, participants could only grab it from the side.

In the real world, the puck’s trajectory was stopped short by a fabric barrier to make it readily available for multiple consecutive slides. However, just before stopping, its velocity was measured, enabling a virtual continuation of the trajectory by simulation. This transition took place after 50cm of the table’s length, which was visually marked by a virtual line for the subjects. To ensure a timely release of the puck, we tracked the subjects’ hand position. After that, the puck’s trajectory was continued via simulation using the basic kinematic equation of motion with constant deceleration (Equation S2). If subjects overreached, or if velocity tracking was faulty, trials had to be repeated. To simplify the problem, we eliminated angles from the simulation, i.e., after passing the measurement line, the puck’s trajectory was directed parallel along the depth of the table Still, to keep the behavior as close as possible to reality, we matched the surface qualities of the table in the simulation. We estimated the friction coefficient between the table’s surface and the puck *c* = 0.29 (and *a* = 9.81*c*) by fitting the deceleration of multiple trajectories recorded with the tracking system.

### General modelling assumptions

Our general approach in modeling was to test multiple models combining different assumptions about motor control, the physical relationships of kinematics, and decision-making in order to test for the hypothesis that participants took the correct physical relationships into account and finding the model best accounting for observed behavior. For an overview of the different models see table 1.

All our models (*Physics, Normal, Linear-Physics*) are instantiations of statistical decision theory, they differ only in their assumptions about variability and utility:

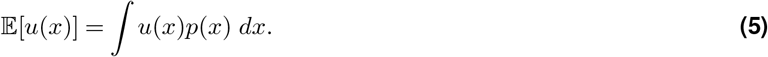

Here, the utility *u*(*x*) captures the costs and benefits of each possible final location *x* of the puck and the term *p*(*x*) of executing a slide with final location *x*. In the plain version of the models *u*(*x*) corresponds to the actual scores *u*(*x*) = *v*(*x*). The models constitute different computational-level descriptions of the task and its optimal solution from the subject’s perspective. From the researcher’s perspective, we build a probabilistic model of the subject’s behavior and estimate the free parameters of the subject’s model using Bayesian methods. Each model includes a free parameter *σ*_*p*_ per participant *p*, which characterizes the variability of movement outcomes. This parameter also determines the optimal action 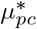 in condition *c* by maximizing expected utility:

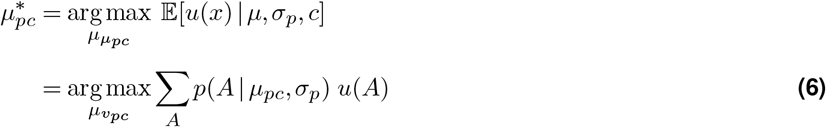

We can discretize expected utility as our possible outcomes consist of separate areas *A* yielding utility *u*(*A*), which are being hit with probability:

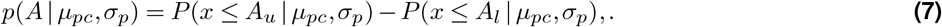

with *A*_*u*_ and *A*_*l*_ being the upper and lower bounds of area *A*.

### The different models

For the *Physics* model, we assume that the generative model of slides fits the task, but also that subjects have access to this model to execute optimal motor actions. The subjects’ task was to generate an initial puck velocity *v*_0_ such that the puck would hit the target but avoid the penalized area (Fig. 1c). This process of generating velocities is subject to signal-dependent variability [43–45] (Fig. 1d). We model this via a log-normal distribution with median *µ* and standard deviation *σ* (see Equation 13):

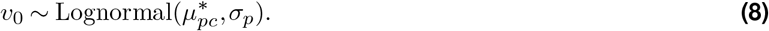

As variability is modeled in the space of velocities, the puck kinematics (Equ. S2) are required to transform velocities *v*_0_ to final puck locations *x*. This implies that 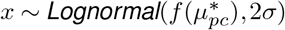, as a random variable which is lognormal distributed and transformed by a quadratic function is again lognormal distributed. The probability of hitting an area *A* for the physics model is then given by:

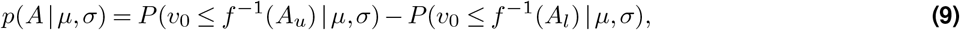

where the inverse *f* ^−1^ yields the velocities need to reach final puck positions.

The *Normal* model embodies the hypothesis that the generative process of sliding can be described with a model that does not account for the increase in variability of final puck position for larger distances, as in [18]. Furthermore, we assume that subjects choose their actions according to the same model for the final puck positions. Therefore, we assume homoscedasticity over the slide outcomes across distances:

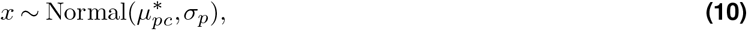

whereby *σ*_*p*_ stays the same across all conditions. Note, that this model does not consider the physical relationship *f*, and instead of considering the problem in the space of initial velocity *v*_*o*_ as the *Physics* model is directly defined in the space of the final puck position *x*.

The *Linear Physics* model is similar to the *Physics* model. The main difference is that it assumes that subjects use a linear heuristic *h* instead of the correct physics to determine the probability of hitting a target:

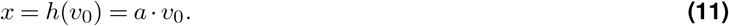

This adds another free parameter *a*_*p*_ to our model for each participant, encoding the assumption that each participant has their own heuristic. From the researcher’s perspective, we are therefore inferring which heuristic the participants would use to account best for the data. From the subject’s perspective, variability in this case only increases with distance because of the signal-dependent noise on the initial puck velocities, but not because of the friction. Note, that in the generative model of the task, the initial velocities and their corresponding slide outcomes are still connected by the correct physical relationship as they would be in the real experiment.

In the plain version of the models, the utility of *u*(*x*) of puck location corresponds to the actual score *u*(*x*) = *v*(*x*). For the subjective utility versions of the models, we included a subjective utility function for negative scores

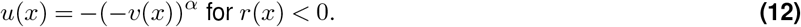

This added another free parameter *α*_*p*_ per participant, modeling the individual subjective utility of receiving penalties.

### Log-normal distribution

We use the log-normal distribution with the following probability density function for modelling:

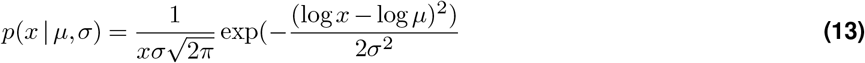

The log-normal distribution inherently models signal-dependent variability [41, 64], as its standard deviation scales linearly with the central tendency: 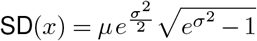

### Model fitting

We implemented the models with the Python probabilistic programming package numpyro [65]. For each model, we drew 4 chains of 1000 warm-up and 10000 samples in using the NUTS-sampler [66] with standard parameters. We made sure that 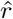 values were close to 1.

#### Efficient sampling of Bayesian actor models

One challenge for implementing our models with probabilistic programming is determining the optimal action *µ*^*^ within the Bayesian actor model, as it involves an optimization problem with no analytic solution. Solving this optimization problem would make sampling of the model highly inefficient. However, one can understand finding the action that maximizes expected utility as a function *g*(*σ*_*p*_, *c*) = *µ*^*^, which depends on the configuration of target and penalty in the experimental condition *c* and on the subject’s motor variability *σ*_*p*_. We computed *g* at each of the distances for a grid of values of penalties and *σ*_*p*_ (penalties: [−10, 0] with step size 0.1, *Physics*: *σ*_*p*_: [0.02, 0.1]; *Normal*: *σ*_*p*_: [0.05, 0.2]; *Linear Physics*: *σ*_*p*_: [0.05, 0.3] with step size 0.001). Consecutively, we fitted these examples with a polynomial of order 10 for each distance, and included the resulting polynomial functions *g*^*′*^ as a replacement for the optimization in the model. This kept the model differentiable, enabling us to use efficient Hamiltonian Monte-Carlo methods [67]. A similar approach would be to approximate *g* with a neural network [51, 68]. As we included a wider range of penalties as the ones used in our experiment, subjective value functions can be easily accommodated for.

#### Prior distributions

As prior for the variability in the space of initial puck velocity *v*_0_, i.e. for the *Physics*, and *Linear Physics* models, we used *σ*_*p*_ ∼ Halfnormal(0, 0.1). Whereas, for the *Normal* model fitting variability in the space of puck final puck location *x*, we used *σ*_*p*_ ∼ Halfnormal(0, 0.2).

#### Linear Models

For the *Linear Model* and the *Subj. Linear Model* optimal aim points are also a function of the heuristic’s slope 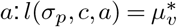. For this function, we used the following grid to fit the polynomial: penalties: [−10, 0] with step size 0.1, *σ*_*p*_: [0.05, 0.3 with step size 0.05 and *a*:[0.05 : 5] with step size 0.1. This allowed as to fit the plain *Linear Model* with the above described MCMC parameters and priors. However, the subjective version of the model proved to be more complicated to fit, as it allows for considerable freedom in the model, while at the same time not being a very good fit to the data. Therefore, we had to resort to using standard random walk metropolis and constraining the priors to exactly the range used to fit the polynomials to approximate *g* with Uniform priors, to make sure the chains do not go out of bounds. This resulted in a reasonable model fit, with 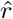 close to 1.

## ACKNOWLEDGEMENTS

This research was supported by ‘The Adaptive Mind’, funded by the Excellence Program of the Hessian Ministry of Higher Education, Science, Research and Art. C.R. acknowledges support by the European Research Council (ERC; Consolidator Award ‘ACTOR’, ERC-CoG-101045783). We would like to thank Nils Neupaertl for useful discussions.

## AUTHOR CONTRIBUTIONS

Conceptualization, Formal Analysis, Visualization: F.T., D.S., C.R.; Investigation, Software, Writing - Original Draft Preparation: F.T.; Supervision, Writing - Review & Editing: D.S., C.R.

## DATA AVAILIABILITY STATEMENT

The source code and data used to produce the results and analyses presented in this manuscript are available from https://osf.io/ck63g/

## COMPETING FINANCIAL INTERESTS

The authors declare no conflicts of interest.

## Supplementary Information

### Methods

#### Kinematic equations

The fundamental kinematic equation

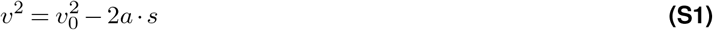

describes how an object’s velocity *v* changes as a function of the initial velocity *v*_0_, the acceleration *a* due to friction, and the object’s displacement *s*. To compute the final puck location *x* (i.e. the final puck displacement) of the hockey puck as a function of its initial velocity, we set the final velocity *v* = 0 and solve for *s*:

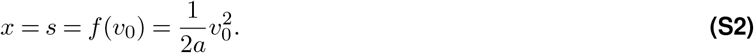

This results in the quadratic function shown in Fig. 1e.

### Statistical Analysis with Mixed Effects Models

#### Hypothesis

1. Variability of initial puck velocity increases with distance.
2. Subjects undershoot more with higher penalty.
3. Subjects undershoot more the larger the target distance.
4. Subjects undershoot more the higher their motor variability.
5. Undershoot is not learned over the course of the trials.

Hypothesis 1 is the only one concerning initial puck velocities *v*_0_. Therefore, we fit a simple regression model with a group level effect of target distance on standard deviation:

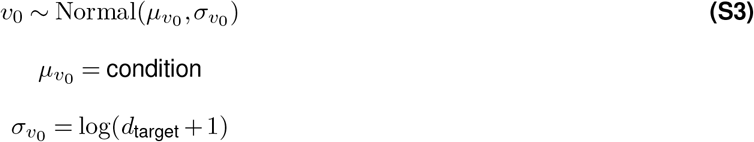

Standard deviation is fit in log space to avoid negative values, but we report the values in the original space for better interpretability.

The other hypotheses were covered in one regression model fitting the deviation of slide endpoints from the target center:

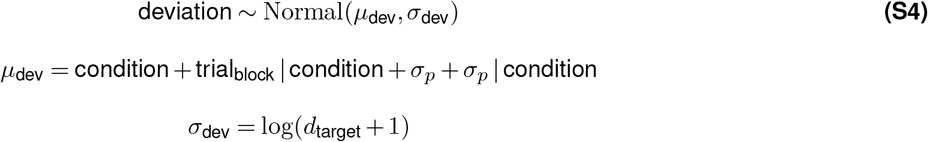

Hypothesis 2 and 3 are included with the effect of condition on *µ*_dev_. We chose this encoding for as straight forward interpretability of interactions between distance in penalty. Therefore, we check for the effect of target distance by comparing the size of condition coefficients across distances, keeping penalty constant. The effect of penalty is tested analogously. Hypothesis 4 is a central prediction of statistical decision theory, and is included by the model by including the per subject variability *σ*_*p*_ as fit by the *Physics* model as a further predictor. Because the model also suggests that the effect of individual variability should be stronger with higher penalty and larger target distance, we additionally include the effect of *σ*_*p*_ per condition. Hypothesis 5 is included by checking whether trial number per block can predict slide end points. Possible learning effects could be different across target condition, therefore we include a group level effect over condition. As a further check for hypothesis 5, we reran the above model on only the first trial of the first session, essentially the first trials participants completed for each target configuration. For this re-run, we excluded the effect of trial_block_. The models were fit with the R package brms [1]. For each model, we drew 4 chains of 1000 warm-up and 10000 samples in using the NUTS-sampler [2] with standard parameters. We made sure that 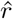 values were close to 1.

### Results

1. We find that standard deviation significantly increases with distance (see SI Fig. S1): 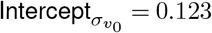, 95% CI [0.120, 0.125] 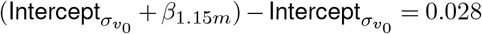, 95% CI [0.025, 0.032] 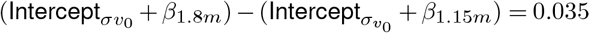, 95% CI [0.030, 0.039]
2. We find that subjects undershoot more with higher penalty: *β*_0,1.8*m*_ − *β*_−1,1.8*m*_ = −0.09, 95% CI [−0.12, −0.07] *β*_−1,1.8*m*_ − *β*_−3,1.8*m*_ = −0.04, 95% CI [−0.07, −0.01] *β*_−3,1.8*m*_ − *β*_−5,1.8*m*_ = −0.05, 95% CI [−0.08, −0.02]
3. We find that subjects undershoot more with target distance: *β*_−5,1.15*m*_ − *β*_−5,0.5*m*_ = −0.06, 95% CI [−0.07, −0.04] *β*_−5,1.15*m*_ − *β*_−5,1.8*m*_ = −0.15, 95% CI [−0.17, −0.12]
4. see main text Results: Subjects’ performance takes into account how their variability interacts with the kinematics.
5. see main text Results: Subjects immediately undershoot more with higher penalty and larger distance. Consider also the results from fitting the model on the first trial per block see Figure. S3.

For an overview over all effect sizes see Figure S2.

## Supplementary Figures

**Figure S1.**
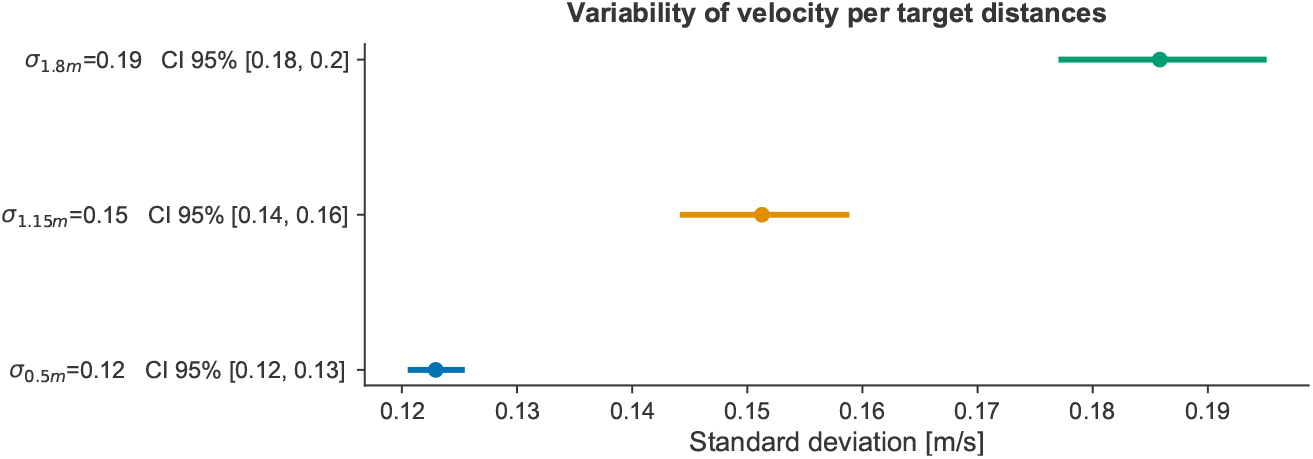
Mixed effects model (Equation SI 1) describing the distance dependent variability of initial puck velocity.

**Figure S2.**
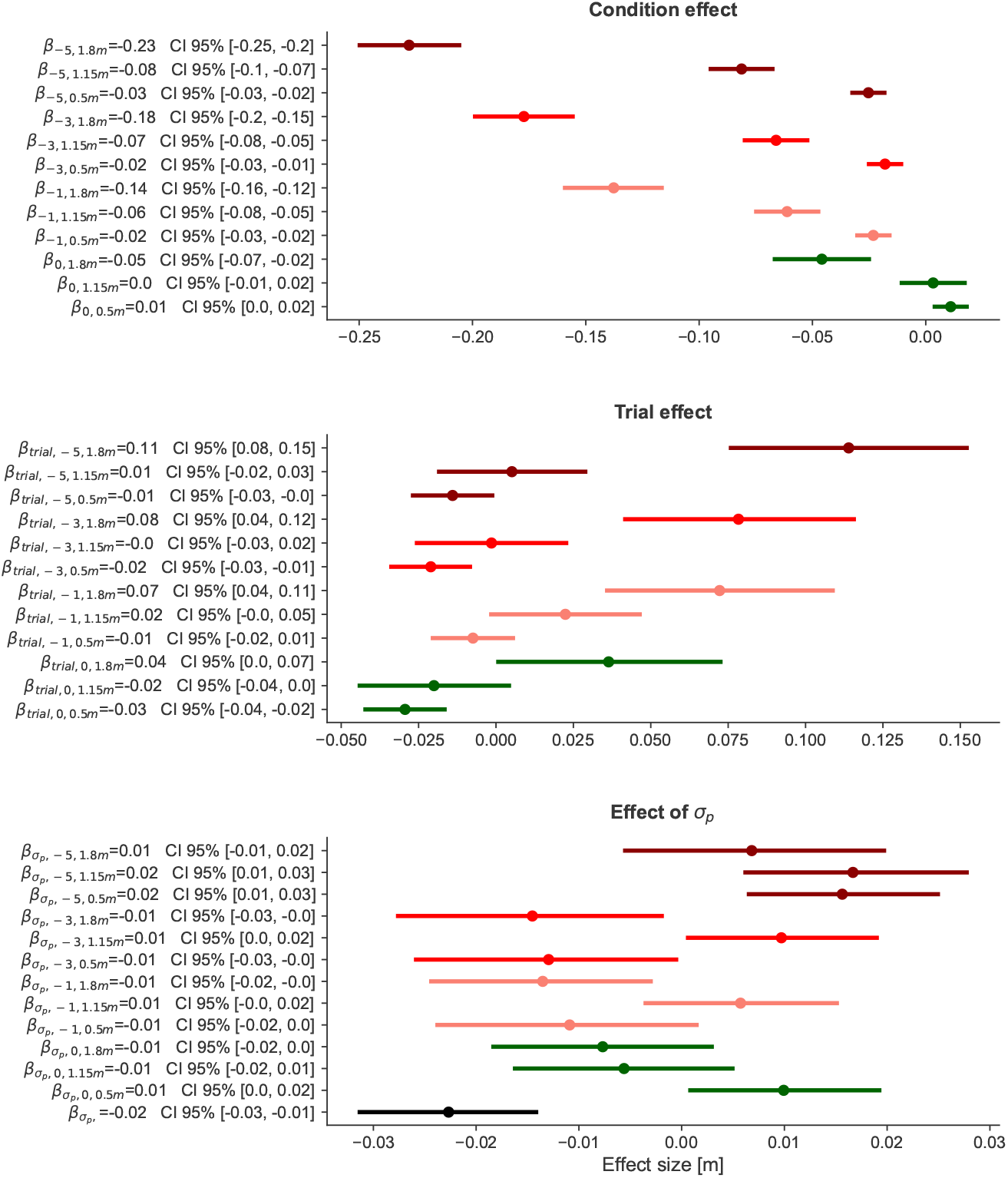
Mixed effects model (Equation SI) describing the influence of condition (categorical coding of target distance and penalty), trial in a block (numerical, normalized between 0 and 1 and individual participant variability *σ*_*p*_ as estimated by the *Physics* model (numerical, z-scored) on the deviation of final puck locations from the target center. Note that we find a significant main effect of *σ*_*p*_ (lowest row) which is slightly lower or higher per condition (other rows).

**Figure S3.**
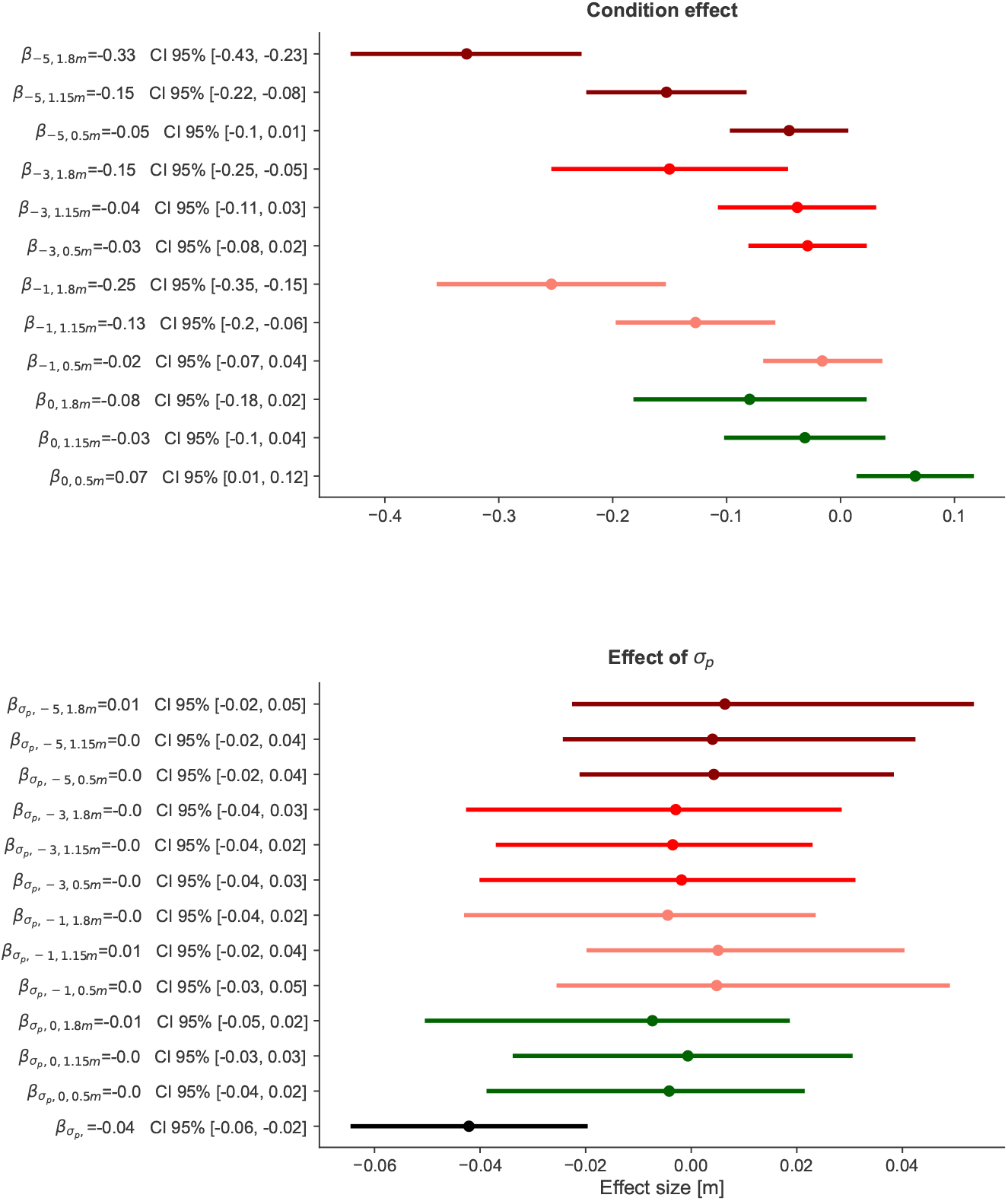
Mixed effects model on the first trials per participant in each condition at the first session (Equation SI without trial_block_) describing the influence of condition (categorical coding of target distance and penalty) and individual participant variability *σ*_*p*_ as estimated by the *Physics* model (numerical, z-scored) on the deviation of final puck locations from the target center. Note that we find a significant main effect of *σ*_*p*_ (lowest row) which is slightly lower or higher per condition (other rows).

**Figure S4.**
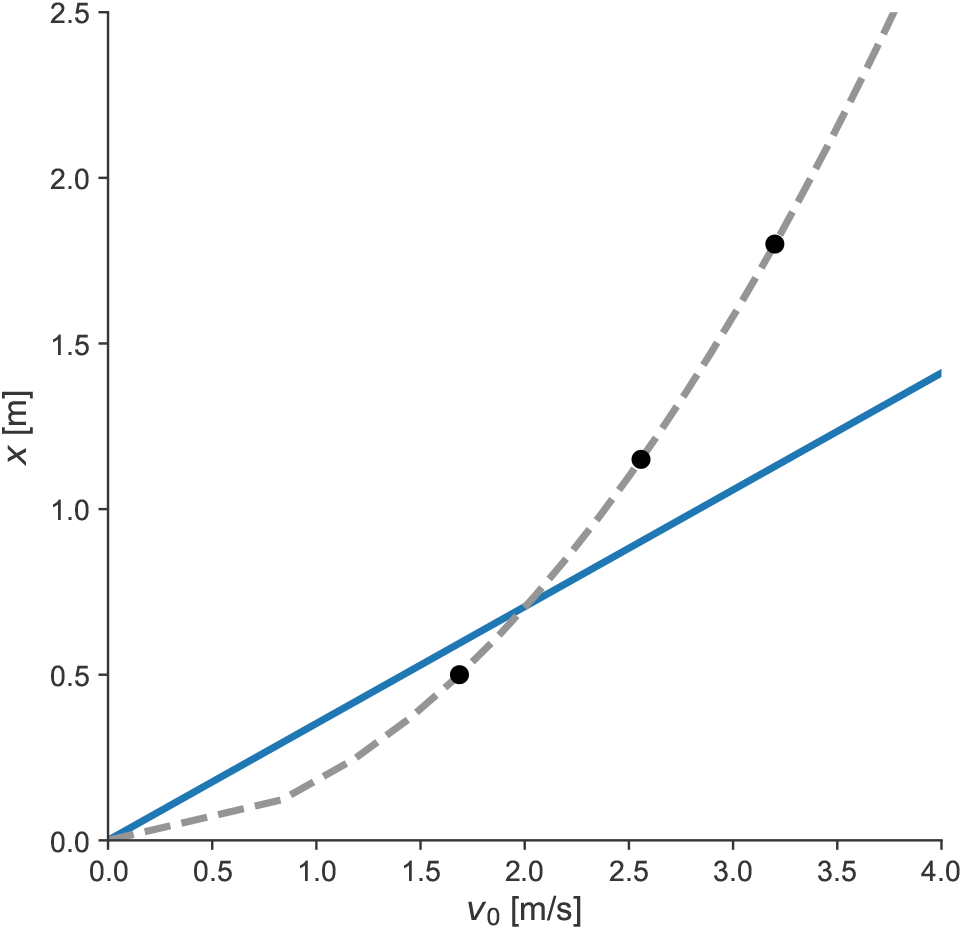
A linear heuristic connecting initial velocity with puck displacement. The three black points correspond to the three target distances used in our task. The linear function has no offset, as a velocity of 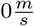 should correspond to 0*m* displacement. The slope of the displayed linear function is the posterior mean across all subjects as fitted with the *Linear* model. Note, the large difference between the linear approximation and the quadratic ground truth for large distances. If a subject assumed the correct velocity needed to hit a far location is determined by a linear function, they would choose a far too large initial puck velocity. Because of the much steeper slope of the quadratic relationship of the real friction, this large velocity would result in an even larger overshoot of the target.

**Figure S5.**
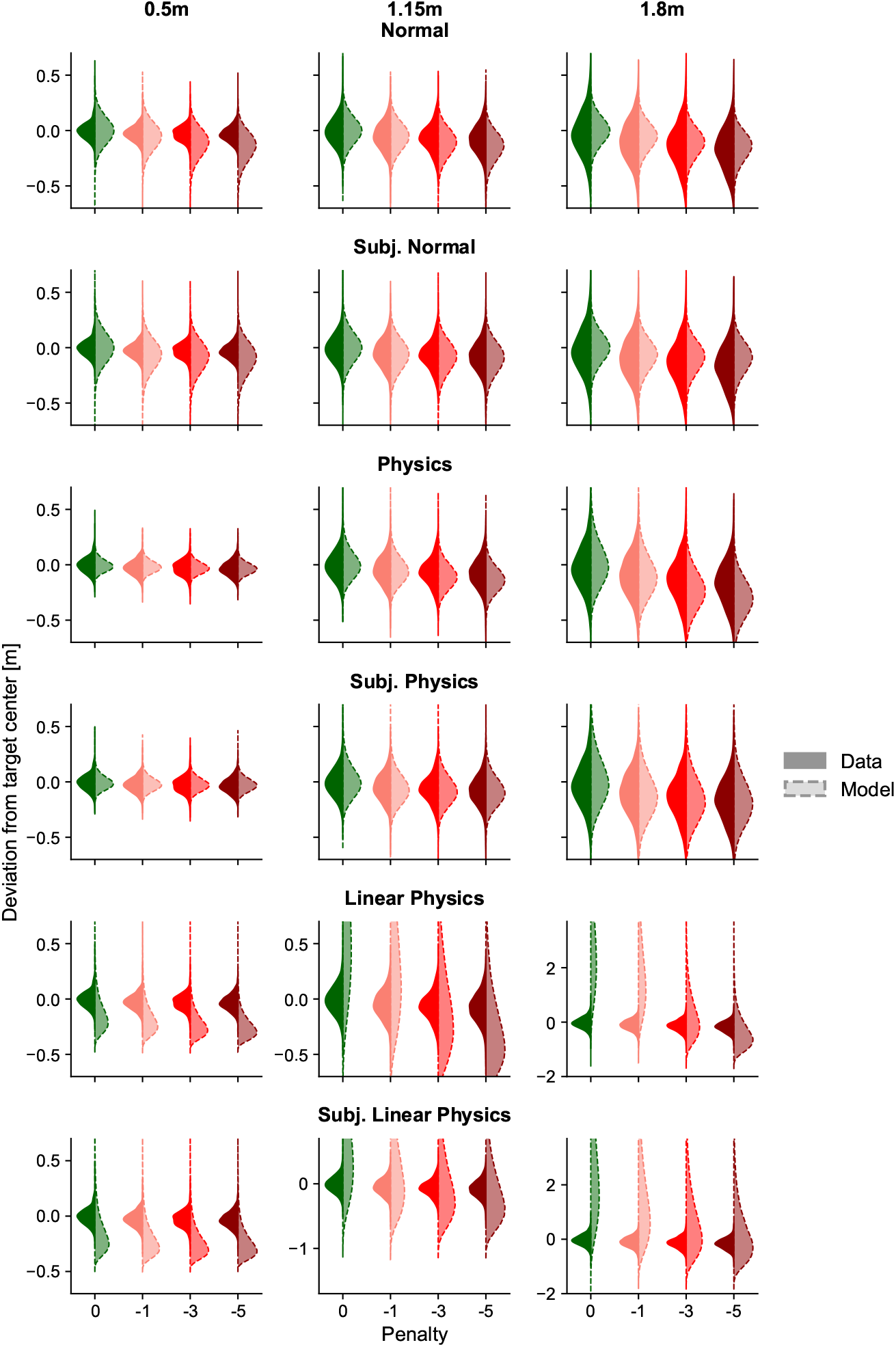
Posterior predictives for all seven models.

**Figure S6.**
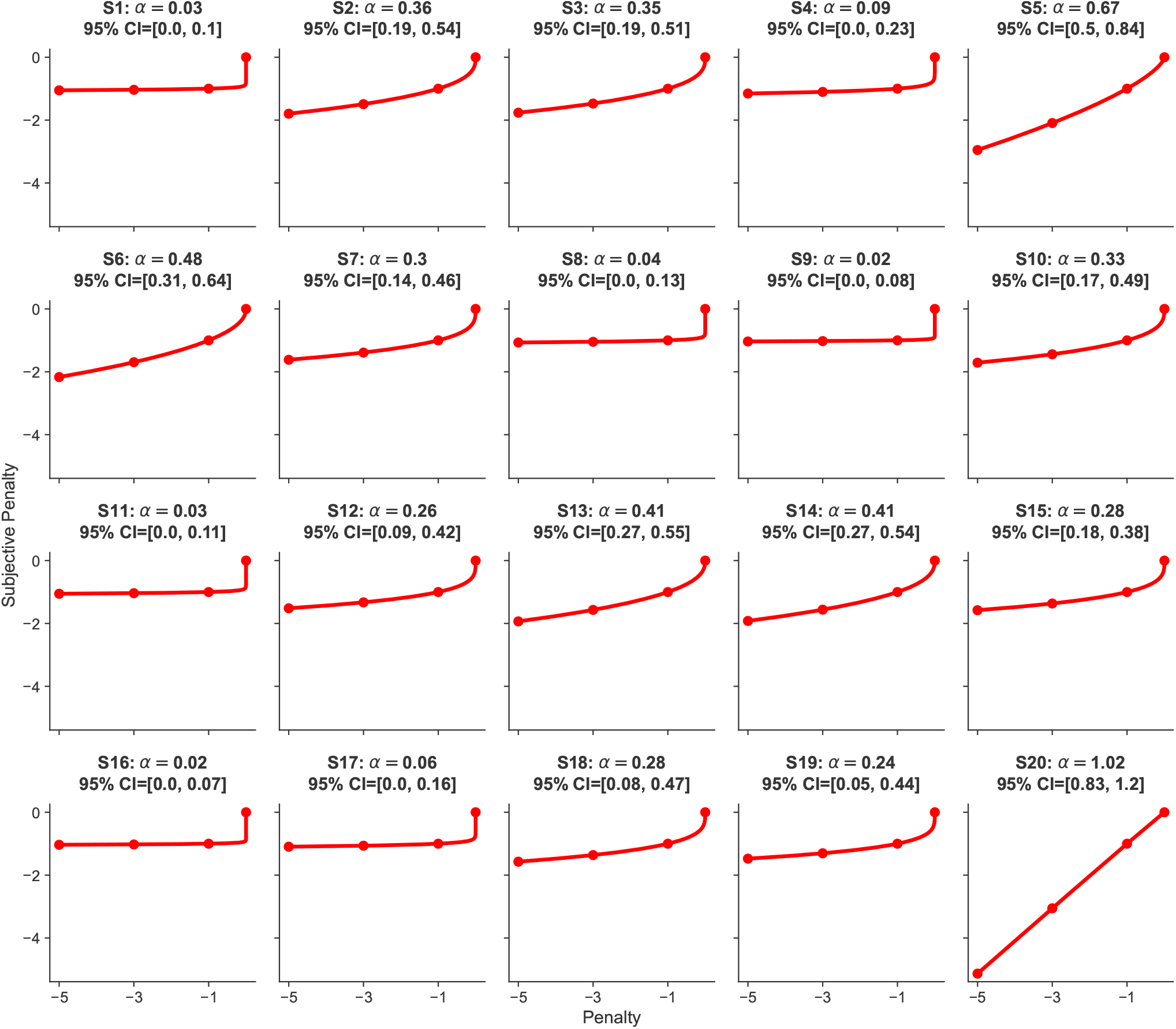
Subjective penalty functions for penalties as fit with the *Subjective Physics* model (posterior mean and confidence intervals of *α*_*p*_)

## Supplementary Material

**Movie S1: Experimental Setup**

## Notes

https://osf.io/ck63g/

### Competing Interest Statement

The authors have declared no competing interest.

### Summary of Updates

Updated the link to the osf repository.

